# Two mutations in the SARS-CoV-2 spike protein and RNA polymerase complex are associated with COVID-19 mortality risk

**DOI:** 10.1101/2020.11.17.386714

**Authors:** Georg Hahn, Chloe M. Wu, Sanghun Lee, Julian Hecker, Sharon M. Lutz, Sebastien Haneuse, Dandi Qiao, Dawn DeMeo, Manish C. Choudhary, Behzad Etemad, Abbas Mohammadi, Elmira Esmaeilzadeh, Michael H. Cho, Rudolph E. Tanzi, Jonathan Z. Li, Adrienne G. Randolph, Nan M. Laird, Scott T. Weiss, Edwin K. Silverman, Katharina Ribbeck, Christoph Lange

**Author notes:** both authors contributed equally (Georg Hahn, Chloe M. Wu).

## Abstract

**Background:** SARS-CoV-2 mortality has been extensively studied in relation to host susceptibility. How sequence variations in the SARS-CoV-2 genome affect pathogenicity is poorly understood. Whole-genome sequencing (WGS) of the virus with death in SARS-CoV-2 patients is one potential method of early identification of highly pathogenic strains to target for containment.

**Methods:** We analyzed 7,548 single stranded RNA-genomes of SARS-CoV-2 patients in the GISAID database (Elbe and Buckland-Merrett, 2017; Shu and McCauley, 2017) and associated variants with reported patient’s health status from COVID-19, i.e. deceased versus non-deceased. We probed each locus of the single stranded RNA of the SARS-CoV-2 virus for direct association with host/patient mortality using a logistic regression.

**Results:** In total, evaluating 29,891 loci of the viral genome for association with patient/host mortality, two loci, at 12,053bp and 25,088bp, achieved genome-wide significance (p-values of 4.09e-09 and 4.41e-23, respectively).

**Conclusions:** Mutations at 25,088bp occur in the S2 subunit of the SARS-CoV-2 spike protein, which plays a key role in viral entry of target host cells. Additionally, mutations at 12,053bp are within the ORF1ab gene, in a region encoding for the protein nsp7, which is necessary to form the RNA polymerase complex responsible for viral replication and transcription. Both mutations altered amino acid coding sequences, potentially imposing structural changes that could enhance viral infectivity and symptom severity, and may be important to consider as targets for therapeutic development. Identification of these highly significant associations, unlikely to occur by chance, may assist with COVID-19 early containment of strains that are potentially highly pathogenic.

---

Viral mutations can cause increased virulence/pathogenicity (Long et al., 2020), both in animals (Geoghegan and Holmes, 2018; Brault et al., 2007), and in humans (Bae et al., 2018; Nogales et al., 2017). Especially for the SARS-CoV-2 virus, the discovery of potential links between viral mutations and disease outcome would have important implications for COVID-19 surveillance and containment (Lo and Jamrozy, 2020), diagnosis, prognosis and treatment development. To identify potential links between viral mutations and mortality, we utilized the GISAID database (Elbe and Buckland-Merrett, 2017; Shu and McCauley, 2017), which currently contains data on 7,548 COVID-19 patients from 86 countries for whom full metadata is available, i.e. age, sex, location and patient status, and whose viral genomes have been sequenced (see Table 1). We probed each locus of the single stranded RNA of the SARS-CoV-2 virus for direct association with host/patient mortality. The variable “patient status” indicates if the patient was alive or deceased at the time the virus sample was submitted to GISAID; we use it as a surrogate for mortality in our analysis. For the analysis, we repurposed the methodology of genome-wide association studies (GWAS) (Manolio, 2010). This approach is widely used in human genetics and can test thousands of genetic loci for association in datasets such as the one of GISAID.

**Table 1:** Characteristics of all patients in the GISAID dataset for whom complete meta-information and sequenced viral genomes were available. Total number of samples (as well as males/females), numbers of deceased/non-deceased, rate of deceased samples at enrollment, mean age, and mutation frequencies for 12,053bp and 25,088bp.

| region | #total | #females | #males | deceased /<br>non-deceased | %deceased | mean age | Mutation frequency in<br>% at the following loci |  |
| --- | --- | --- | --- | --- | --- | --- | --- | --- |
|  |  |  |  |  |  |  | 12,053 | 25,088 |
| entire dataset | 7548 | 3313 | 4235 | 722 / 6826 | 9.6 | 47.6 | 1.2 | 2.2 |
| Africa | 1517 | 954 | 563 | 2 / 1515 | 0.1 | 38.8 | 0.0 | 0.2 |
| Eastern Mediterranean | 730 | 180 | 550 | 131 / 599 | 17.9 | 45.4 | 0.0 | 0.1 |
| Europe | 1872 | 896 | 976 | 70 / 1802 | 3.7 | 56.0 | 0.1 | 0.0 |
| Pan American Health Organization | 1505 | 637 | 868 | 435 / 1070 | 28.9 | 51.9 | 5.7 | 10.6 |
| Brazil | 430 | 223 | 207 | 192 / 238 | 44.7 | 55.1 | 20.0 | 37.0 |
| South-East Asia | 1116 | 367 | 749 | 83 / 1033 | 7.4 | 45.1 | 0.0 | 0.1 |
| Western Pacific | 808 | 279 | 529 | 1 / 807 | 0.1 | 41.6 | 0.0 | 0.2 |

To identify potential confounding geographic factors in the sequencing data, we first conducted principal component analysis of the Jaccard similarity matrix (Figure 1) that was computed for the 7,548 viral genomes available for our analysis. We utilized the Jaccard similarity matrix because its computation does not require estimates of the mutation frequency for each locus in the SARS-CoV-2 genome, in contrast to other similarity matrices such as the variance/covariance matrix (Prokopenko et al., 2016). We found that the virus genomes clustered in distinctive branches that correspond to the geographic regions from where their data was submitted to GISAID (Forster et al., 2020, Hahn et al., 2020), see Figure 1. The geographical clustering of the viral genomes can cause bias in the association analysis if unaccounted for. Hence, we generated additional eigenvector plots to investigate the number of eigenvectors needed to eliminate bias caused by such clustering. Based on visual inspection of these plots, we selected the first 10 eigenvectors of the Jaccard matrix as covariates for the following logistic regression analyses.

**Figure 1:**
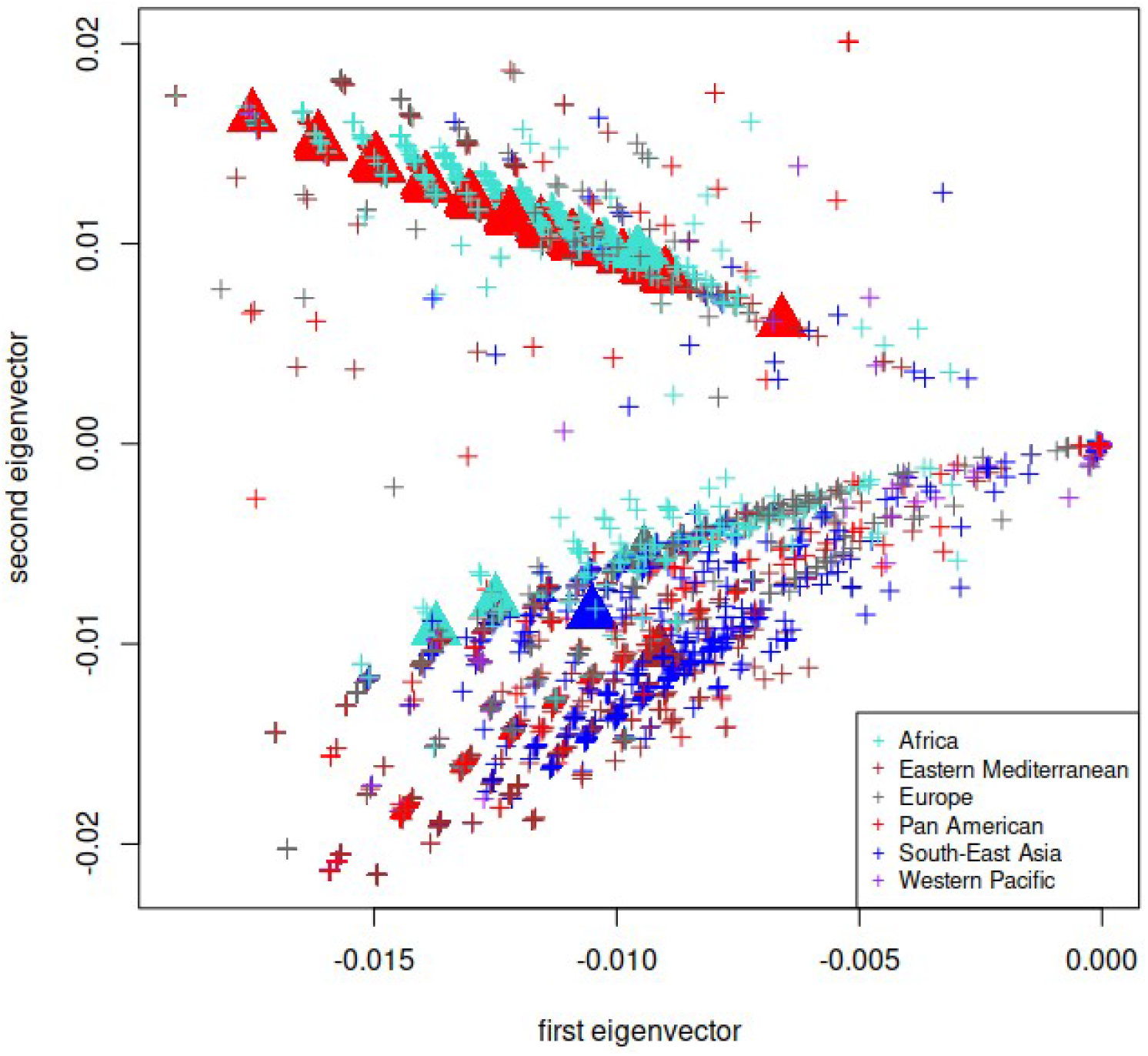
Geographic distribution of 7,548 SARS-CoV-2 genomes. Genomes are depicted according to their first two eigenvectors of the Jaccard matrix and colored by geographic region. The eigenvector plot shows distinct grouping of SARS-CoV-2 genomes according to their geographic origin. Furthermore, genomes that carry a mutation at 12,053bp or 25,088bp are depicted by triangles. The majority of those are located in a subbranch whose samples come predominantly from Pan America.

## Methods

### Data acquisition

The analysis presented in this article is based on nucleotide sequences with accession numbers EPI_ISL_403962 to EPI_ISL_636981, downloaded from the GISAID database (Elbe and Buckland-Merrett, 2017; Shu and McCauley, 2017) as a file in “fasta” format on 15 November 2020. Only patients with additional metadata (age, sex, and hospitalization status as plain text comments) were selected on GISAID, resulting in 8,647 samples.

### Data cleaning

We filtered the 8,647 samples for complete nucleotide sequences, and aligned them to the SARS-CoV-2 reference sequence (published on GISAID under the accession number EPI_ISL_402124) using MAFFT (Katoh et al., 2002).

Using the location tag in the fasta file, we grouped all samples according to the WHO regional offices for Africa (AFRO), for the Eastern Mediterranean (EMRO), for Europe (EURO), for South-East Asia (SEARO), for the Western Pacific (WPRO), as well as the Pan American Health Organization (PAHO). In particular, the countries included in each group are as follows: (1) AFRO (Algeria, South Africa, Gambia, Nigeria, Senegal, as well as Congo, Madagascar, Mozambique, Tunisia, Ghana, Rwanda, Cameroon); (2) EMRO (Egypt, Morocco, Kuwait, Lebanon, Oman, Saudi Arabia, United Arab Emirates, as well as Iran, Iraq, Bahrain); (3) EURO (Austria, Belgium, Bosnia and Herzegovina, Bulgaria, Croatia, Cyprus, Czech Republic, Denmark, Faroe Islands, France, Germany, Hungary, Italy, Israel, Poland, Portugal, Romania, Russia, Slovakia, Spain, Sweden, Turkey, Kazakhstan, as well as Andorra, Georgia, Norway, Ukraine, Switzerland, Saint Barthelemy, Guadeloupe, Saint Martin, Mongolia, Greece, Finland, Moldova, Reunion); (4) PAHO (Canada, USA, Costa Rica, Mexico, Argentina, Brazil, Chile, Colombia, Ecuador, Peru, Venezuela, as well as Puerto Rico, Uruguay, Panama, Dominican Republic); (5) SEARO (Bangladesh, India, Indonesia, Myanmar, Nepal, Sri Lanka, Thailand); (6) WPRO (Cambodia, Japan, Malaysia, Vietnam, Australia, Guam, Hong Kong, China, Singapore, as well as South Korea, Taiwan, New Zealand, Philippines).

Finally, we matched the samples to the metadata information (age, sex, clinical outcome) available on GISAID. Filtering for those samples having complete metadata information resulted in n=7,548 samples.

### Data analysis

After alignment with MAFFT (Katoh et al., 2002), we compared all aligned sequences of length p=29,891 entrywise to the SARS-CoV-2 reference sequence, and denoted in a matrix X with an entry X_ij_=1 that sequence i deviated from the reference sequence at position j. All other entries of X are zero.

We used the R-package “locStra” (Hahn et al., 2020c,d) to calculate the Jaccard similarity matrix (Jaccard, 1901; Tan et al., 2005; Prokopenko et al., 2016; Schlauch et al., 2017) for the n viral genomes based on the matrix X. The Jaccard matrix J(X) has n rows and n columns, and each entry (i,j) is the Jaccard similarity index between the i’th and j’th SARS-CoV-2 genome in our dataset. Computation of the first 10 eigenvectors of the Jaccard similarity matrix J(X) allows us to visualize the geographic clustering of the viral genomes. We also guard the logistic regression analysis against confounding by including the first eigenvectors in the regression analysis as covariates.

For the association analysis of the entire viral genome, we defined the response to be a binary indicator for the clinical outcome, where we only distinguish between all those patients/hosts whose hospitalization status tag at enrollment into the GISAID database was listed as “deceased” (outcome of 1) versus the remaining samples as non-deceased (outcome of 0). At this point, no other information regarding clinical outcome is available in GISAID.

We performed a logistic regression of the binary outcome variable for each of the p=29,891 loci on the following covariates: the column vector X_·i_ encoding the mismatches of each sample at the i’th location on the SARS-CoV-2 nucleotide sequence, the patient’s age, sex, location (WHO region), and the first 10 eigenvectors of the Jaccard matrix. The WHO region was included as we observed in Figure 1 that the viral genomes cluster into distinct branches that correspond to the geographic regions. The logistic regression was carried out in R using the default “glm” command, where the parameter “family” was set to “family=binomial(link=“logit”)”. We tested the i’th locus/location of the viral genome for association with mortality by testing whether the regression coefficient for column X_·i_ is equal to zero. We controlled for multiple tests using the Bonferroni correction at an uncorrected threshold of 0.05, resulting in the corrected threshold of 0.05/29,891=1.67e-06.

Finally, we also perform an analysis with a matched dataset. For this, we match each sample in GISAID that is deceased at submission to the closest non-deceased one, measured in Euclidean distance in the eigenvector space of the Jaccard-matrix (Figure 1). When running the logistic regression on the matched dataset, we test each of the p=29,891 loci on the column vector X_·i_ (encoding the mismatches to the reference genome), as well as the patient’s age and sex only.

## Results

### Whole-genome association analysis of the SARS-CoV-2 genomes

After testing each locus (presence/absence of mutation) of the viral genome individually for association with the status indicator variable (deceased/non-deceased) of the host/patient at submission to GISAID, two loci of the SARS-CoV-2 genome achieved genome-wide significance: one at position 12,053bp with p-value 4.09e-09, and one at 25,088bp with p-value 4.41e-23 (Table 2).

**Table 2:** Sample size, number of deceased samples, as well as p-values and odds ratios from the logistic regression on the two mutations: for the entire dataset, for each WHO region, and for samples from Brazil only.

| analysis | sample size | deceased | locus | p-value | odds ratio |
| --- | --- | --- | --- | --- | --- |
| overall | 7548 | 722 | 12,053 | 4.09e-09 | 6.4 |
|  |  |  | 25,088 | 4.41e-23 | 12.9 |
| Matched analysis | 1452 | 722 | 12,053 | 5.53e-05 | 3.5 |
|  |  |  | 25,088 | 4.91e-11 | 4.8 |
| PAHO | 1505 | 435 | 12,053 | 1.22e-09 | 7.3 |
|  |  |  | 25,088 | 3.10e-24 | 15.9 |
| Brazil | 430 | 192 | 12,053 | 2.27e-04 | 3.5 |
|  |  |  | 25,088 | 4.90e-13 | 9.2 |

To investigate the robustness of the highly significant association signals, we examined the dataset at the individual patient and locus level. Our findings were enabled by two features specific to the data: 1.) the Brazilian centers enrolled much larger numbers of deceased patients than the other centers world-wide. At enrollment, 44.7% of the Brazilian patients were deceased in contrast to only 9.6% in the entire dataset. 2.) We also noticed that all genomes that carry at least one of the mutations either at 12,053bp or 25,088bp are located predominantly in the branch of the eigenvector plot (see Figure 1) that corresponds to the PAHO/South America region.

We conducted two different types of sensitivity analyses to minimize the chances that the observed associations are caused by confounding/GISAID dataset composition (Table 2): 1. Our data set was restricted to genomes that were matched based proximity in the eigenvector plots (see the Methods section for details), called “matching” in Table 2. 2. As further examination of the deceased indictor variable revealed that all “deceased” carrier genomes came from Brazil, our second sensitivity analysis was restricted to genomes that were submitted from the PAHO region and Brazil, respectively. In both analyses, 25,088bp maintained significance at 0.05/29,891=1.67e-06, and 12,053bp stayed borderline significant. both loci, the effect size estimates of the mutations showed risk increases for mortality of a factor of 3.5-7 for carriers of a mutation at 12,053bp, and a factor of 5-16 for carriers of a mutation at 25,088 (Table 2).

To summarize, all results of the secondary analyses (Table 2) support the genome-wide significant association between the mutation 25,088bp and mortality. The locus at 12,053bp did not formally achieve genome-wide significance in the secondary analyses (when matching and restricting the analysis to Brazil only), but nonetheless remains a viable candidate locus. The large effect estimates for both mutations (Table 2) are substantial in support of the associations. Since the criteria for selection into the study likely varies by country, and may be related to the deceased indicator, the odds ratio estimate from the Brazil sample alone may be most interpretable. Among the samples from Brazil, 18.2% of the patients whose viral genome did not carry any mutation at either loci were deceased at enrollment, compared with 82.4% for patients whose viral genomes carried the mutation at 25,088bp only, and 82.6% for those carrying a mutation at both 12,053bp and 25,088bp.

Given the large effect estimates for mutations in all analyses (Table 2), it is difficult to imagine an unaccounted confounding mechanism that would affect mutations at just two out of almost thirty-thousand loci and that would be strong enough to cause such profound association signals, as the ones we observed in our analysis. Table 1 also provides a regional breakdown of the “deceased-at-enrollment” rates and the mutation frequencies for both loci. The rarity of the mutations outside of Brazil means that there is virtually no power to detect any association (if they exist).

## Discussion

Single mutations in viruses can confer enhanced virulence associated with patient mortality (Bae et al., 2018; Brault et al., 2007). In our analysis of SARS-CoV-2, the mutation at 25,088bp occurs in the spike glycoprotein, which mediates viral attachment and cellular entry. The spike protein consists of two functional subunits: S1, which contains the receptor-binding domain, and S2, which contains the machinery needed to fuse the viral membrane to the host cellular membrane. The mutation at 25,088bp is in the S2 subunit, and specifically occurs within the S2’ site, which is cleaved by host proteases to activate membrane fusion (Figure 2). In many viruses, membrane fusion is activated by proteolytic cleavage, an event which has been closely linked to infectivity—for instance, a multibasic cleavage site is a signature of highly pathogenic viruses including avian influenza (Walls et al., 2020). In coronaviruses, membrane fusion is known to depend on proteolytic cleavage at multiple sites, including the S1/S2 site, located at the interface between the S1 and S2 domains, and the S2’ site located within the S2 domain. These cleavage events can impact infection—in fact, a distinct furin cleavage site present in the SARS-CoV-2 S1/S2 site is not found in SARS-CoV (Vankadari, 2020), and it is thought to increase infectivity through enhanced membrane fusion activity (Walls et al., 2020; Vankadari, 2020; Xia et al., 2020). Consequently, mutations at these sites can alter virulence—for instance, a recent study reported that mutations disrupting the multibasic nature of the S1/S2 site affect SARS-CoV-2 membrane fusion and entry into human lung cells (Hoffmann et al., 2020). Several studies have also found that SARS-CoV mutants with an added furin recognition site at S2’ had increased membrane fusion activity (Belouzard et al., 2009; Watanabe et al., 2008). While enhanced infectivity does not always cause a higher fatality rate, more infectious viruses can lead to a higher viral load, which can impact symptom severity and mortality (Pujadas et al., 2020).

**Figure 2:**
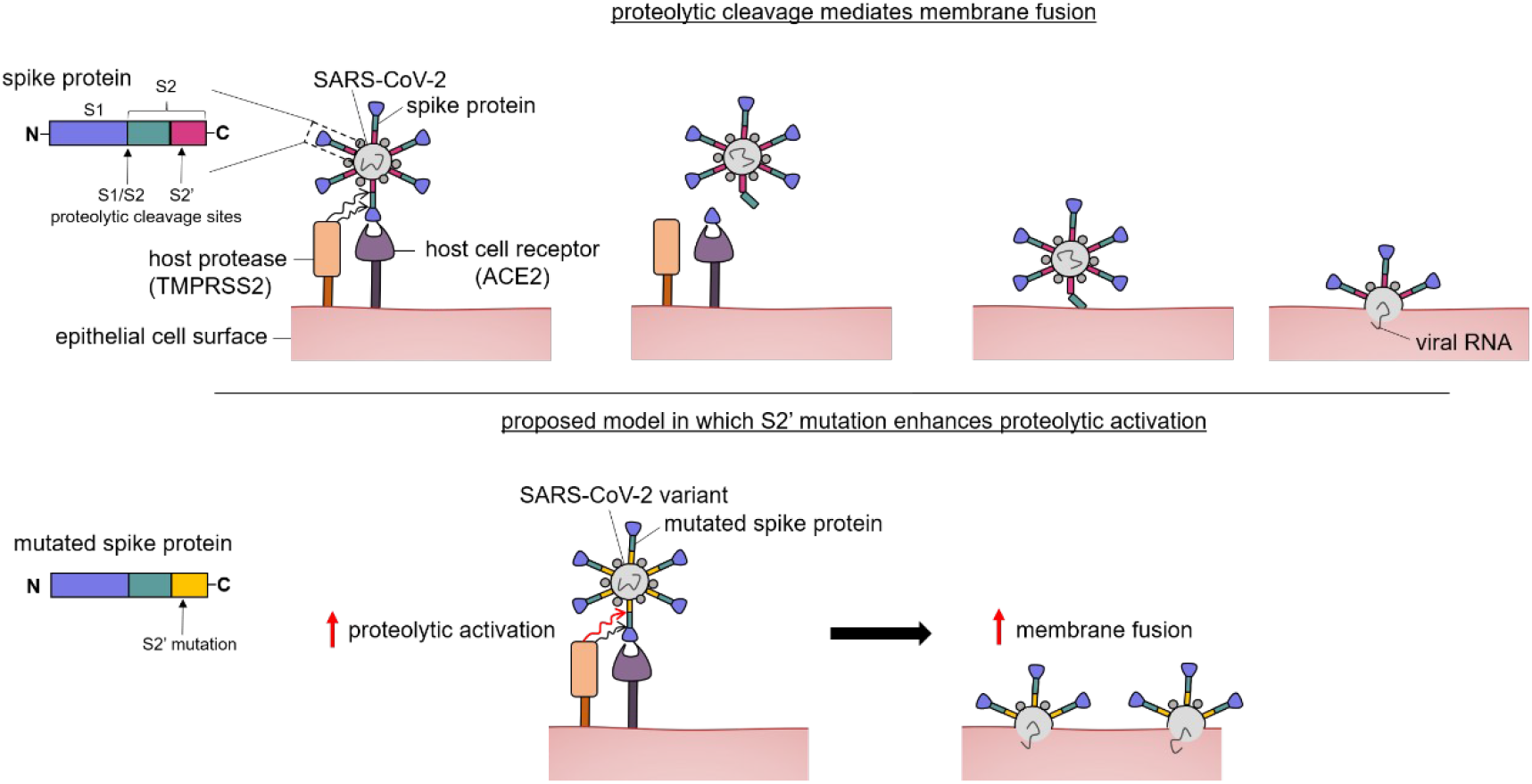
Proposed model showing how the S2 mutation may enhance proteolytic activation. The SARS-CoV-2 spike protein is colored by region (blue—S1, green—S2, magenta—S2’). The S2’ site is cleaved by host proteases, facilitating membrane fusion and viral entry into host cells. A mutation in this region, depicted in yellow, could theoretically increase proteolytic activity and membrane fusion, thereby causing greater infectivity.

All carriers of a mutation at 25,088bp exhibit a G to T missense mutation (Table 3), which changes the encoded amino acid from valine to phenylalanine. Compared to the branched-chain structure of valine, phenylalanine has a bulkier aromatic structure. Such a substitution may impose local structural constraints, stabilize particular secondary structures (Makwana and Mahalakshmi, 2015), or introduce specific interactions which lead to preferential binding. Therefore, a mutation in the S2’ domain which promotes proteolytic cleavage could theoretically enhance viral infectivity (Figure 2) and consequently, patient mortality. While many current therapies primarily target the receptor binding domain within the S1 subunit of the SARS-CoV-2 spike protein, our findings suggest that the S2 domain may be an important additional target for therapeutic development.

**Table 3:** Number of genomic variants at each locus, affected protein position, and corresponding amino acid change. Amino acid in the reference sequence in bold.

| Locus | A | C | G | T | Protein | Position | Primary substitution |
| --- | --- | --- | --- | --- | --- | --- | --- |
| 12,053 | 0 | **7453** | 0 | 87 | nsp7 | 71 | Leu --> Phe |
| 25,088 | 0 | 0 | **7331** | 166 | Spike | 1176 | Val --> Phe |

The mutation at 12,053bp occurs within the ORF1ab gene, which expresses a polyprotein comprised of 16 nonstructural proteins (Yoshimoto, 2020). Specifically, 12,053bp occurs in NSP7, which dimerizes with NSP8 to form a heterodimer that complexes with NSP12, ultimately forming the RNA polymerase complex essential for genome replication and transcription. Mutations causing enhanced viral polymerase activity have been linked to increased pathogenicity of influenza viruses. All carriers of a mutation at 12,053bp exhibit a C to T missense mutation, which causes leucine to be substituted for phenylalanine (Table 3). Such a mutation may confer structural rigidity which could potentially alter interactions with other components of replication and transcription machinery, but experimental analysis is needed to test these hypotheses.

Collectively, these results suggest that genetic variation in the viral genome sequence may contribute to the increased COVID-19 mortality. Although biological follow-up experiments are needed for functional validation, early containment of highly pathogenic viral strains during a pandemic may require early intervention when biostatistical extreme associations are identified.

## Acknowledgements

The authors gratefully acknowledge the contributors, originating and submitting laboratories of the sequences from GISAID’s EpiCoV™ Database (Elbe and Buckland-Merrett, 2017; Shu and McCauley, 2017) on which this research is based. A detailed list of contributors is available in the Supplementary Information.

## Data Availability Statement

Sequence data that support the findings of this study are deposited in the GISAID database with accession numbers in the range of EPI_ISL_403962 to EPI_ISL_636981 (https://www.gisaid.org/).

